# Midbody inheritance predicts re-entry into quiescence, but not lineage potential, in mouse hematopoietic stem cells

**DOI:** 10.64898/2026.07.28.739739

**Authors:** Tsuyoshi Fukushima, Arne Wehling, Ren Shimamoto, Shuhei Asada, Shunsuke Kawamura, Tomofusa Fukuyama, Susumu Goyama, Timm Schroeder, Toshio Kitamura, Yosuke Tanaka

## Abstract

Hematopoietic stem cells (HSCs) give rise to all blood cell lineages and possess long-term self-renewal potential. HSCs undergo symmetric division for their expansion and asymmetric division to generate one HSCs and one progenitor cells which contribute to production of mature blood cells. The midbody is a structure which is formed in the center of the intercellular bridge during cytokinesis. However, the midbody is either asymmetrically inherited by one daughter cell or symmetrically released after cell division, whether these distinct patterns of midbody inheritance influence HSC fate remain poorly understood. In this study, we designed a fusion protein hmKO2 and MgcRacGAP which is a component of midbody. We then traced the midbody inheritance during cell division and the future cell fates of HSC daughters after division by time-lapse imaging. As a result, we found that the midbody release correlated with the delay of the time to the next division but not to the lineage potential of HSCs, indicating the possibility that midbody remnant plays some roles in cell cycle progression.

**Highlight:** Hematopoietic stem cells exhibit a low frequency of midbody inheritance.

Midbody inheritance does not affect the lineage potential of daughter cells.

Midbody loss is associated with delayed entry into the next cell cycle.

## Introduction

Hematopoietic stem cells (HSCs) give rise to all blood cell lineages and possess self-renewal potential[1]. HSCs undergo symmetric division to expand the stem cell pool and asymmetric division to generate one HSC and one progenitor cell, thereby contributing to the production of mature blood cells. The balance between symmetric and asymmetric division is thought to regulate both HSC pool size and mature blood cell production.[2-5]. Several proteins and organelles have been reported to be asymmetrically inherited by daughter cells. In HSCs, Numb, mitochondria, and lysosomes have been reported to be asymmetrically inherited, and their relationships with HSC fate have been investigated.[2, 3, 6, 7]. However, the relationship between the asymmetric inheritance of protein and organelle and the cell fate of HSCs is not completely clear.

The midbody is a structure formed at the center of the intercellular bridge during cytokinesis[8, 9]. The midbody consists of the central spindle, the contractile ring [8, 9], the actomyosin contractile ring[8, 9], and proteins including MgcRacGAP. MgcRacGAP is a GTPase-activating protein (GAP) which plays pivotal roles in actin ring contraction and completion of cytokinesis [10-12]. At the end of cytokinesis, one side of the intercellular bridge is cleaved and the midbody is asymmetrically inherited by one daughter cell, or both sides of the intercellular bridges are cleaved and midbody is released from both cells[13]. Thus, the midbody is either asymmetrically inherited or symmetrically released and not inherited by either daughter. However, the relationship between midbody inheritance and the cell fate of HSCs is still poorly understood.

In this study, we generated an MgcRacGAP–hmKO2 fusion protein to visualize the midbody during HSC division. Using time-lapse imaging, we tracked midbody inheritance during cell division, the subsequent lineage potential of the HSC daughters, and the time to the next division. We found that midbody inheritance was not associated with the lineage potential of daughter HSCs. In contrast, midbody release was associated with a longer time to the next division, suggesting that midbody inheritance may predict HSC re-entry into quiescence.

## Material and methods

### Mice

Female C57BL/6 were purchased from Japan SLC (Shizuoka, Japan). Eight-to twelve-week-old male and female mice served for experiments unless otherwise noted. All mice were housed at 22 ±2 □ and 12:12 hrs light: dark cycle. The experiments were approved by the Committee on the Ethics of Animal Experiments, University of Tokyo.

### Cell lines

PlatE[14] was cultured in D-MEM (Wako) containing 10% fetal bovine serum (FBS) (biowest). PlatE was authenticated by short-tandem repeat analyses and tested for mycoplasma contamination in our laboratory. All cell lines were cultured at 37 □ and 5 % CO2.

### Antibodies, Flow Cytometry and Cell Purification

Hematopoietic cells were isolated from femurs and tibias, humerus, sternum, pelvic and vertebra by clashing and depleted of red blood cells by ACK Lysing Buffer (Thermo Fisher Scientific). Cells were stained with the appropriate dilution of fluorochrome- and/or biotin-conjugated antibodies and Propidium iodide (PI) for dead cell exclusion and analyzed on FACS Aria (BD Biosciences). The following antibodies were used: fluorescent CD150 (clone TC15-12F12.2), CD48 (clone HAM48-1), CD34 (clone RAM34), c-Kit (clone 2B8), Sca-1 (clone D7) and biotinylated lineage antibodies (CD11b, Gr-1, TER119, CD45R/B220, CD5) (BioLegend). The sorted cells were collected into ice-cold PBS.

### Viral Transduction

Retroviruses were produced in Plat-E cells cultured in 10-cm dishes by transient transfection using the calcium phosphate method as previously described [15, 16]. Viral supernatants were collected 48 h after transfection, concentrated by centrifugation (15,000 × g, 4 h), and resuspended in 50 μl of StemSpan SFEM (STEMCELL Technologies). LT-HSCs were sorted from C57BL/6 mice and cultured for 12 h before infection. RetroNectin-coated 96-well plates were prepared by overnight incubation with 100 μl of 25 μg/ml RetroNectin (TaKaRa Bio) at 4°C, followed by blocking with 2% BSA. LT-HSCs and retrovirus were added to the plates, and time-lapse imaging was performed 24 h after viral transduction.

### Time-lapse imaging

Time-lapse experiments were conducted at 37°C, 20% O2, 5% CO2 on μ-slide VI0,4 channel slides (IBIDI), coated with 10 μg/ml anti-CD43–biotin, in phenol-red-free IMDM supplemented with 20% BIT, 100 ng/ml SCF, 100 ng/ml TPO, 10ng/ml IL3, 4 U/ml EPO, 2 mM l-glutamine, 50 μM 2-mercaptoethanol, 50 U/ml penicillin and 50 μg/ml streptomycin using a Nikon-Ti Eclipse equipped with linear-encoded motorized stage, Orca Flash 4.0 V2 (Hamamatsu), and Spectra X fluorescent light source (Lumencor). White light emitted by the Spectra X was collimated and used as a transmitted light for bright-field illumination via a custom-made motorized mirror controlled by an Arduino UNO Rev3 (Arduino). Fluorescent images were acquired using optimized filter sets: FITC (470/40; 495LP; 525/50), mKO2 (546/10; 560LP; 577/25), Cy5 (620/60; 660LP; 700/75; all AHF) to detect Alexa Fluor 488, hmKO2, APC, respectively. Time intervals of bright-field and fluorescent image acquisition were chosen to minimize photo-toxicity. Images were acquired using 10× CFI Plan Apochromat λ objective (NA 0.45) and 40× CFI Plan Apoλ (NA 0.95). Single-cell tracking and image quantification were performed using self-written software as previously described[2, 17-22].

### Quantification and statistical analysis

Statistical analyses were performed by the unpaired and two-tailed Student’s t-test after testing for normal distribution and equal variance. GraphPad Prism 9 was used for these statistical analyses. All data are presented as mean ± SEM in the figures. Significance levels were set at p* < 0.05, and p** < 0.01.

### Data availability

Information and requests for reagents may be directed to the lead contact Yosuke Tanaka.

## Results and Discussion

To examine midbody inheritance and the subsequent fate of HSCs after cell division, sorted HSCs were transduced with a retroviral vector expressing the MgcRacGAP–hmKO2 fusion protein, and time-lapse imaging was initiated 24 hours after transduction (Figure 1A). The MgcRacGAP–hmKO2 fusion protein labeled both the midbody and the nucleus after S phase (Figure 1B). Daughter cells inheriting the midbody were designated as the inherit group, whereas their sister cells were designated as the non-inherit group (Figure 1B, left). Cell divisions in which the midbody was released from both daughter cells were designated as the release group (Figure 1B, right). The lineage potential of daughter HSCs was determined based on CD71 and CD16/32 expression together with cellular morphology. CD71-positive cells were classified as erythroid progenitors (Er), CD16/32-positive cells as granulocyte/monocyte progenitors (GM) (Figure 1C), and large cells as megakaryocyte progenitors (Mk) (Figure 1D).Time-lapse imaging enabled reconstruction of lineage trees (Figure 1E), allowing evaluation of midbody inheritance, lineage potential, and future cell-cycle progression after cell division (Figure 1E).

**Figure 1.**
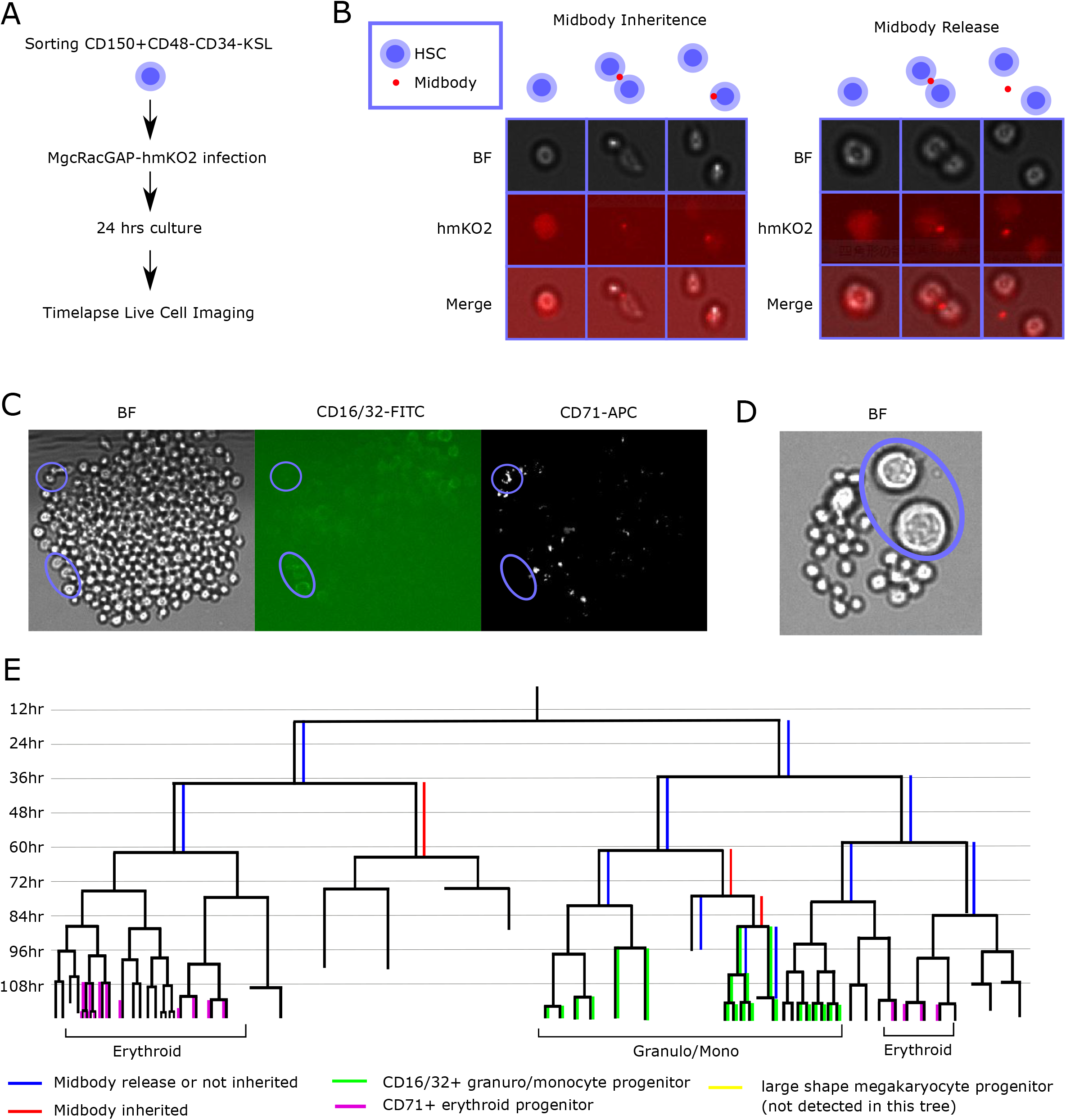
(A) schematic diagram of experiment. (B) Representative picture of midbody inheritance (left) and release (right). Bight field image (BF), hmKO2 and Merged image was shown. Before division, during cytokinesis and after division was shown as left to right. (C) Representative picture for evaluate lineage selection. Bight field image (BF), CD16/32-FITC and CD71-APC was shown. Upper circle contained cells which is CD71+ erythrocyte progenitor. Bottom circle contained cells which is CD16/32+ glanuro/monocyte progenitor. (D) Representative picture for lineage detection by morphology. Bright field image (BF) was shown. Circle contained cells which is large megakaryocyte progenitor. (E) Representative tree diagram drowns from tracking single cells in time-lapse imaging data. Blue line indicated midbody release or not inherited. Both daughters showing blue line indicated midbody release. Red line indicated midbody inheritance. Green line indicated CD16/32+, Purple line indicated CD71+ and Yellow line indicated large Megakaryocyte.

First, we quantified examined midbody inheritance during HSC division. It is known that normal stem cells show higher release rate than cancer cells [13]. In agreement with these observations, the midbody-release rate was high at 76.5% in first division of HSCs (Figure 2A). This rate decreased to less than 50% after the second division (Figure 2A). Because HSCs progressively lose self-renewal capacity, multilineage potential, and quiescence under ex vivo culture conditions [23, 24], we hypothesized that these stem cell properties might be associated with midbody inheritance. However, no correlation was observed between the lineage potential of the mother HSC (i.e. the lineage output of the whole colony) and midbody inheritance (Figure 2B).

**Figure 2.**
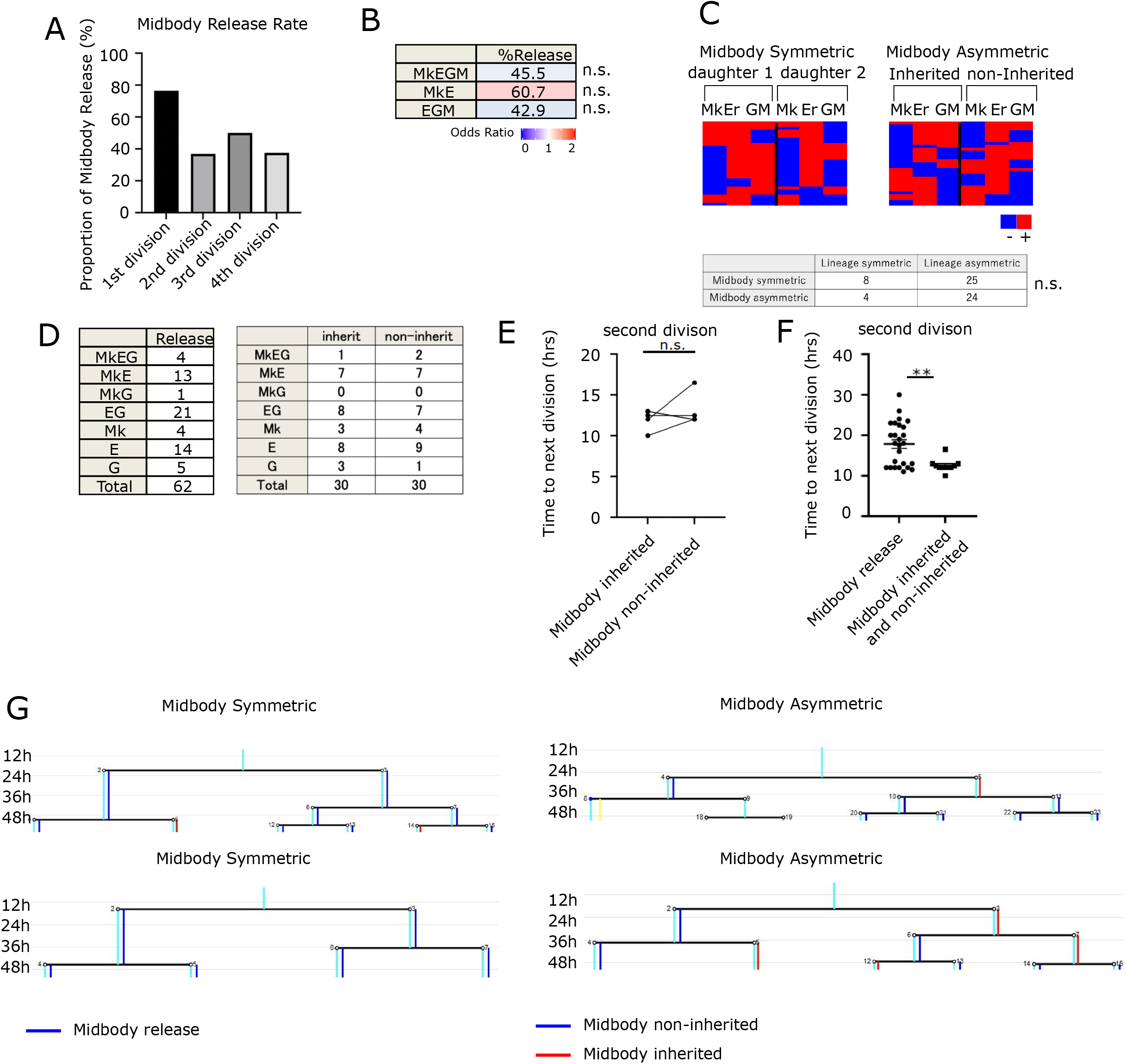
(A) Frequency of midbody release in successive divisions. (B) Frequency of midbody release rate by the mother lineage potential (i.e. the lineage output of the whole colony). Colored by odds ratio. (C) Heatmap indicated lineage selection of daughter cells. Red: lineage detected, Blue: lineage not detected. (Midbody symmetric: left, Midbody asymmetric: right)(upper). `Table showing that` midbody symmetry and lineage symmetry. The association between midbody symmetry and lineage symmetry was assessed using Fisher’s exact test. (bottom) (D) The numbers of clones which showed indicated lineage potential by the inheritance of midbody (Midbody symmetric: left, Midbody asymmetric: right). (E) Time to the next cell division in paired daughter cells following asymmetric midbody inheritance. The time to the next division was compared between the daughter cell that inherited the midbody and its sister cell that did not. Paired t-test. (F) Time to the next cell division in daughter cells from HSC pairs exhibiting symmetric midbody release (release group) or asymmetric midbody inheritance (inherit group and non-inherit group). (G) Representative tree diagram drowns from time-lapse imaging data. Cyan lines represent the cell lineage tree and are shown for visualization only. Blue line indicated midbody release or not inherited. Both daughters showed blue line indicated midbody release. Red line indicated midbody inheritance.

Next, we examined the relationship between HSC daughter midbody inheritance and their individual lineage potential. No bias in lineage potential was observed between sister cells that inherited the midbody, did not inherit the midbody, or were generated following midbody release (Figure 2C, D). Despite the asymmetric inheritance of the midbody, no difference in lineage potential was detected between paired HSC daughter cells, suggesting that midbody inheritance alone is insufficient to alter HCS lineage potential. Previous studies have reported that the midbody can be released into the extracellular space and subsequently internalized by neighboring cells through endocytosis [13]. In the present study, we were unable to distinguish between daughter cells that directly inherited the midbody after asymmetric cleavage of the intercellular bridge and those that acquired the midbody through secondary endocytosis after its release. Distinguishing these two modes of midbody acquisition in future studies may provide further insight into the relationship between midbody inheritance and HSC fate.

Finally, we examined the relationship between midbody inheritance and the time to the next cell division. No difference in the time to the next division was observed between sister cells that inherited the midbody and those that did not (Figure 2E and 2G). However, both daughter cells generated by divisions with midbody release exhibited a significantly longer time to the next division (Figure 2F). These findings indicate that midbody release is associated with delayed entry into the next cell division. Because daughter cells that did not inherit the midbody after asymmetric inheritance did not exhibit delayed cell-cycle re-entry, the midbody itself is unlikely to be directly responsible for this delay. Rather, midbody behavior may serve as a marker, rather than a direct effector, of the cellular state associated with delayed cell-cycle re-entry. Previous studies have shown that the proliferation–quiescence decision is established during the preceding M phase [25]. Because midbody inheritance is a cytokinetic event that occurs during M phase, our observation that midbody release is associated with delayed entry into the next cell cycle raises the possibility that midbody release reflects cellular events associated with re-entry into quiescence after division.

## Acknowledgments

We thank the Flow Cytometry Core at The Institute of Medical Science, The University of Tokyo for their help. This project was supported by ‘‘Grant-in-Aid for Challenging Exploratory Research’’ from JSPS Grant Number 17K19645 to Y.T. and Grant Number 20K21613 to Tsu.F.

